# Effects of Release of TSG-6 from Heparin Hydrogels on Supraspinatus Muscle Regeneration

**DOI:** 10.1101/2024.08.20.608812

**Authors:** Joseph J. Pearson, Jiahui Mao, Johnna S. Temenoff

**Author notes:** Corresponding Author: Johnna S. Temenoff Address: 315 Ferst Dr. NW Atlanta, Georgia 30332, Author: Joseph J. Pearson, Address: 315 Ferst Dr. NW Atlanta, Georgia 30332, Author: Jiahui Mao, Address: 1660 Peachtree ST NW APT 5305, Atlanta, Georgia 30309.

## Abstract

Muscle degeneration after rotator cuff tendon tear is a significant clinical problem. In these experiments, we developed a poly(ethylene glycol)-based injectable granular hydrogel containing two heparin derivatives (fully sulfated (Hep) and fully desulfated (Hep-)) as well as a matrix metalloproteinase-sensitive peptide to promote sustained release of Tumor Necrosis Factor Stimulated Gene 6 (TSG-6) over 14+ days *in vivo* in a rat model of rotator cuff muscle injury. The hydrogel formulations demonstrated similar release profiles *in vivo*, thus facilitating comparisons between delivery from heparin derivatives on level of tissue repair in two different areas of muscle (near the myotendious junction (MTJ) and in the muscle belly (MB)) that have been shown previously to have differing responses to rotator cuff tendon injury. We hypothesized that sustained delivery of TSG-6 would enhance the anti-inflammatory response following rotator cuff injury through macrophage polarization, and that release from a fully sulfated heparin derivative (Hep) would potentiate this effect throughout the muscle. Inflammatory/immune cells, satellite cells, and fibroadipogenic progenitor cells, were analyzed by flow cytometery 3 and 7 days after injury and hydrogel injection, while metrics of muscle healing were examined via immunohistochemistry up to Day 14. Results showed controlled delivery of TSG-6 from Hep caused heightened macrophage response (Day 14 macrophages, 4.00 ± 1.85% single cells, M2a, 3.27 ± 1.95% single cells) and increased markers of early muscle regeneration (embryonic heavy chain staining) by Day 7, particularly in the MTJ region of the muscle, compared to release from desulfated heparin hydrogels. This work provides a novel strategy for localized, controlled delivery of TSG-6 to enhance muscle healing after rotator cuff tear.

**IMPACT STATEMENT:** Rotator cuff tear is a significant problem that can cause muscle degeneration. In this study, a hydrogel particle system was developed for sustained release of an anti-inflammatory protein, Tumor Necrosis Factor Stimulated Gene 6 (TSG-6), to injured muscle. Release of the protein from a fully sulfated heparin hydrogel-based carrier demonstrated greater changes in amount inflammatory cells and more early regenerative effects than a less-sulfated carrier. Thus, this work provides a novel strategy for localized, controlled delivery of an anti-inflammatory protein to enhance muscle healing after rotator cuff tear.

## 1. INTRODUCTION

The prevalence of degenerative rotator cuff disease increases with age, often leading to tendon tear.^1,2^ Tendon tear can result in chronic muscle degeneration including inflammation, atrophy, fatty infiltration, fibrosis and overall fiber disorganization.^3^ Moreover, it has been found that rotator cuff muscle has a limited ability to regenerate, even after tendon reattachment^4^. Thus, therapies targeting resolution of the chronic inflammatory state of the muscle could provide improved outcomes for these patients.

A hallmark of muscle degeneration after rotator cuff injury is the persistence of inflammatory cells, including monocytes, macrophages, and dendritic cells^5^, as well as elevated matrix remodeling enzymes such as matrix metalloproteinases (MMPs)^6^. In previous work, our laboratory has shown that MMP2 is significantly increased 2 weeks after rotator cuff injury in a rat model^7^. Thus, a potential means to affect the composition of the local muscle environment after rotator cuff tear is local delivery of soluble anti-inflammatory signals that can reduce MMP activity.

In particular, Tumor Necrosis Factor Stimulated Gene 6 (TSG-6) is a broadly anti-inflammatory protein that can address multiple aspects of the degenerative muscle environment. TSG-6 has inherent immunomodulatory properties that can aid in polarization of macrophages away from pro-inflammatory (M1) and towards pro-healing phenotypes (M2)^8^, as well as inhibit plasmin that is the upstream regulator of MMPs^9^. Additionally, injection and/or cell secretion of TSG-6 has been shown to decrease pro-inflammatory cytokines while increasing anti-inflammatory cytokines in mouse models for colitis, hepatitis and lung injury.^8,10–13^

In previous work, it has been found that the effects of TSG-6 can be enhanced by complexing with glycosaminoglycans (GAGs)^14^. GAGs are a group of negatively-charged natural polymers that include hyaluronic acid, heparin, heparan sulfate, and chondroitin sulfate^15^. The negative charge of GAGs allows for electrostatic interaction with positively-charged growth factors such as TSG-6^16^. Among other interactions, TSG-6 can complex with heparin to potentiate inter-alpha inhibition of plasmin^9,17^. This has led our laboratory and others to explore complexation with GAGs to reduce tissue damage in models of osteoarthritis^18,19^. Our laboratory found that removing sulfation from heparin reduced the anti-plasmin activity of the complex. In particular, fully sulfated heparin (Hep) potentiated the anti-plasmin effects of TSG-6 compared to desulfated heparin (Hep-).^18^

GAG-containing hydrogels and hydrogel microparticles have been extensively explored to deliver positively-charged growth factors such as stromal derived factor 1α^4^, bone morphogenetic protein 2^20^, as well as TSG-6^18^. However, it can be difficult to control the placement of *in vivo* injection due to hydrogel gelation time and diffusion of microparticles in a dynamic environment. In response, various laboratories have created granular hydrogels, which consist of previously-crosslinked microgels with interactions between the microgels that provide structural support.^21^ Since these are already crosslinked structures, the concern of hydrogel gelation time is removed and the structural support creates a largely consistent placement of the material and thereby consistent therapeutic release.

Another important aspect of protein delivery vehicles is their potential for degradation *in vivo*^22^. Given the elevated MMP activity after rotator cuff tear^23^, we focused our attention on inclusion of enzyme-sensitive peptide sequences to promote local delivery at the site of muscle injury. Specifically, our laboratory has developed an MMP responsive polyethylene glycol (PEG)-based hydrogel system^24^ that was adopted for use in this work.

Thus, through these studies, we have developed a novel fragmented PEG-based hydrogel system including heparin and MMP-sensitive peptides for loading and delivery of TSG-6. We hypothesized that sustained delivery of TSG-6 from a fragmented hydrogel system would enhance and prolong the anti-inflammatory response following rotator cuff injury through macrophage polarization, and that release from Hep would potentiate this TSG-6 effect. To examine this hypothesis, three groups were investigated in a rat rotator cuff injury model: Saline injection (injured control), Hep + TSG-6 (Hep) and Hep-+ TSG-6 (Hep-). Muscle degeneration and myogenic response were examined via immunohistochemistry and enzyme linked immunosorbent assays (ELISAs) for MMP activity. Macrophage and T-cell response was examined via flow cytometry. Flow cytometry was also performed for cells that are known to respond to TSG-6 (neutrophils, mesenchymal stem cells (MSCs))^8^ and muscle-resident cells (satellite cells (SCs) and fibroadipogenic progenitor cells (FAPs)).

## 2. MATERIALS AND METHODS

### 2.1 MMP Cleavable PEG Synthesis

MMP cleavable PEG materials were synthesized using previously published techniques.^24^ Briefly, acryl-PEG-SVA (3.4kDA, Laysan Bio) was conjugated with the peptide, GGVPMSMRGGGK, at a 1:2.2 molar ratio, respectively. The conjugation was conducted in 50mM sodium bicarbonate buffer (pH 8.5) for 4 hours. The solution was then dialyzed (3.5 MW cutoff, Spectrum Labs) for 48 hours with regular water changes. After dialysis, the solution was aliquoted in scintillation vials, frozen in liquid nitrogen, lyophilized (Labconco) and stored at −20°C.

### 2.2 Heparin Biomaterial Synthesis

Hep-was synthesized similarly to previous methods.^18^ Briefly, 10 mg/mL of heparin sodium salt (Hep) from porcine intestinal mucosa (Sigma) was dissolved in distilled water and passed through Dowex 50WX4 resin (Sigma). The heparin solution was combined with pyridine (Sigma) to reach a pH of 6. The heparin/pyridine solution was placed in a rotary evaporator (Buchi) to remove excess liquid and then was flash frozen and lyophilized. The heparin pyridium salt was added to a dimethyl sulfoxide (DMSO, VWR)/distilled water (9:1 v/v) solution at 1 g/mL and reacted at 100°C for 24 hours, at which point the heparin solution was precipitated in an ethanol (Decon Laboratories)/sodium acetate solution (VWR). The precipitate was collected via centrifugation, dissolved in distilled water, dialyzed, lyophilized and stored at −20°C.

Hep and Hep-were then functionalized with methacrylamide (Mam, Polyscience)) resulting in HepMAm and Hep-MAm as previously described^25,26^. Briefly, to functionalize methacrylamide onto Hep and Hep-, 40 mg of disodium phosphate (Fisher) and 4.5 g of monosodium phosphate (Fisher) were dissolved in 100 mL of distilled water to make a buffer solution. Then, 200 mg of Hep or Hep-, 180 mg of N-Hydroxysulfosuccinimide sodium salt (sulfo-NHS, Sigma) and 180 mg of N-(3-Aminopropyl)methacrylamide hydrochloride (APMAm, Polyscience) were mixed in a scintillation vial. 10 mL of the buffer solution was added to the scintillation vial and stirred to dissolve. 150 mg of N-(3-Dimethylaminopropyl)-N’-ehtylcarbodiimide hydrochloride (EDC, Sigma) was added to the solution and stirred to dissolve on ice for 2 hours. After 2 hours, another 150 mg of EDC was added to the solution and stirred to dissolve on ice for 4 hours. The solution was aliquoted, lyophilized for 4 days and stored at - 20°C.

### 2.3 Hydrogel Fragment Fabrication and Characterization

The MMP cleavable PEG material was dissolved in sterile saline. For fragments containing HepMAm or Hep-MAm, the heparin derivative was added to the PEG solution and incubated at 37°C for 30 minutes. Ammonium persulfate (APS) and tetramethylethylenediamine (TEMED) were added to a final concentration of 35 mM. The solution was mixed and added to a 1mL syringe (NormJect) with attached, sealed needle and was placed in an incubator (HERA Cell 150) at 37°C for 30 minutes for crosslinking. The hydrogel was then extruded from the syringe through a 40 µm filter (PluriSelect) into 1.5 mL centrifuge tubes (100 µL/tube) to initialize fragmentation. Deionized water was then added into the tubes. The fragment solutions were mixed and passed through the 40 µm filter a total of 4 times. The fragments were then flash frozen in liquid nitrogen, lyophilized and frozen at −20°C. For sterile fragmentation, all solutions were sterilized through a 0.22 µm filter, fragmentation was conducted in a biosafety cabinet on a sterile field, and a 0.22 µm filter (Corning) was used during lyophilization.

For characterization, 3 batches of fragments were synthesized and 100 µL of fragment solution were added to a glass slide and imaged (Nikon Eclipse TE2000-U, 4x) and analyzed with Image J (NIH). Since the fragments are irregular in shape, two length measurements were taken for at least 300 fragments per batch accounting for the longest and shortest length of each fragment. The average length per fragment was calculated and the batches were compared (n=3 batches, at least 300 fragments per batch).

### 2.4 Rotator Cuff Injury Model

Rotator cuff injury was induced as previously described.^4,25^ Eight to ten week old male Sprague-Dawley rats (initially 250-300 g) were ordered and utilized in accordance with protocols approved by Georgia Institute of Technology Institutional Animal Care and Use Committee. 5% isoflurane was administered initially to induce anesthesia and then maintained between 2-3% during the procedure. The surgical area was shaved and sanitized with alcohol and chlorhexidine (Povidone-iodine was used in lieu of chlorhexidine for infrared imaging). A ∼2 cm incision was made parallel to and slightly below the left clavicle through skin and the deltoid muscle. This incision provided access to resect a ∼0.5 mm segment of the suprascapular nerve to induce denervation. Using the window created by the incision through the deltoid, both the supraspinatus and infraspinatus tendons were transected. To reduce the potential for spontaneous reattachment, PharMed BPT tubing (Tygon, 1.6 mm inner diameter) was secured to the end of each tendon via 4-0 sild non-absorbable sutures (Oasis). The deltoid muscle and skin were subsequently closed with Vicryl 4-0 sutures (Ethicon) and wound clips, respectively. Sustained release buprenorphine was administered as an analgesic.

### 2.5 *In vivo* Growth Factor Release

For *in vivo* growth factor release, TSG-6 (R&D Systems) was fluorescently tagged with Alexa Fluor 647 NHS ester (Invitrogen, Microscale Kit) by combining the TSG-6 and the Alexa Fluor 647 NHS ester in 100mM sodium bicarbonate buffer, incubating at room temperature to 15 minutes and then centrifuging with the kit’s resin to obtain the final tagged TSG-6 Fluorescently tagged TSG-6 was combined with either Hep or Hep-containing fragments at molar ratios of 6.7:1 and 666.7:1, respectively, in sterile saline to yield a 100 mg/mL solution. The solutions were placed on a tube rotator (ThermoScientific) overnight at 4°C to allow for TSG-6 loading.

For *in vivo* assessment of the delivery platforms, Hep + TSG-6 or Hep-+ TSG-6 systems were injected into the supraspinatus muscle of rats after rotator cuff injury procedure using a 22g needle (BD) inserted on the edge of the left scapula proximal to the shoulder slightly above the spine of the scapula. The needle was inserted parallel to the spine of the scapula for 10 mm at which point the syringe plunger was depressed while simultaneously removing the needle from the muscle leading to a linear delivery of 50 µL of fragment solution to the muscle. Using an *In Vivo* Imaging System (IVIS, Perkin Elmer), fluorescent images of the injection region were taken 1, 3, 7, 14 and 21 days after injection measuring the average radiant efficiency with reduced values over time equating to release of the AF647 tagged TSG-6 (n=3-4 animals per group).

### 2.6 *In vivo* Assessment of TSG-6 Treatments

For *in vivo* assessment of the delivery platforms, Hep, Hep-, or sterile saline was injected (20 µL) into the supraspinatus muscle as previously mentioned in the IVIS methods. The Hep and Hep-groups contained 1.6:1 and 166.7:1 molar ratio to TSG-6, respectively, delivering 1 µg of TSG-6 per injection. These groups were evaluated using flow cytometry (n=5-8 animals per group) and immunohistochemistry detailed below (n=3-5 animals per group).

### 2.7 Flow Cytometry

Flow cytometry (Cytek Aurora) was completed for harvested supraspinatus muscles to assess changes in inflammatory cells and subtypes, muscle specific cells and mesenchymal stem cells (BioLegend, BD, Bio-Rad, Bioss, antibodies found in Table 1).^4,25,27–32^ The supraspinatus muscles were prepared for flow cytometry by harvesting, splitting the muscle into two regions (first 10 mm from the musculotendinous junction distally along the muscle (MTJ) and the remainder of the muscle belly (MB)), and mincing. The contralateral supraspinatus muscles were used as controls. All samples were digested in collagenase type II (Worthington), Dispase II (ThermoFisher), and DMEM (ThermoFisher) solution at 37°C. The solution was diluted, the digested muscles were filtered (Fisher, 22363547), centrifuged (Thermo) and resuspended in 3% fetal bovine serum (FBS, ThermoFisher) in phosphate buffered saline (PBS, Corning). The samples were split for the flow panel and stained in the dark for 30 minutes. The samples were diluted, centrifuged and fixed with 2% paraformaldehyde (PFA, J.T. Baker). Lastly, the samples were diluted to 200 µl in PBS, centrifuged and resuspended with PBS at stored at 4°C until flow cytometry acquisition.^4^ The resulting data was gated (FlowJo) and analyzed using a two-way ANOVA with posthoc Tukey’s test (significant difference, p<0.05) and with Spanning-tree Progression Analysis of Density-normalized Events (SPADE) clustering to identify cell population changes^33^.

**Table 1:** Flow cytometry cell populations.

| Cell Type | Markers |
| --- | --- |
| **Leukocytes** | CD11b+ |
| **Neutrophils** | CD11b+CD45+RP-1+ |
| **M1 Macrophage** | CD11b+CD68+CD86+ |
| **M2c Macrophage** | CD11b+CD68+CD163+ |
| **M2a Macrophage** | CD11b+CD68+CD206+ |
| **T-cell** | CD3 |
| **Cytotoxic T-cell** | CD3+CD8+ |
| **Helper T-cell** | CD3+CD4+ |
| **Regulatory T-cell** | CD3+CD4+CD25+ |
| **MSC (endogenous)** | CD45-CD31-CD90+CD29+CD44+ |
| **Satellite cells** | CD45-CD31-Sca1-VCAM-1+ |
| **Fibro-adipogenic progenitor cells** | CD45-CD31-Sca1+VCAM-1- |

### 2.8 Immunohistochemistry (IHC) and Image Analysis

7 and 14 days after rotator cuff injury and treatment, the injured and contralateral supraspinatus muscles were harvested and embedded in 100% optimum cutting temperature (OCT, Sakura Finetek) following consecutive incubation in 10% v/v OCT/PBS, OCT/PBS with 10% sucrose (VWR), and OCT/PBS with 20% sucrose under vacuum. The 100% OCT containing muscles were then placed under vacuum overnight followed by freezing in hexanes (Sigma) solution surrounded by liquid nitrogen cooled ethanol (Fisher Scientific) solution. The embedded muscles were sectioned at 10 µm via cryostat (Thermo Scientific CryoStar NX70) with sections collected in both the MTJ and MB regions as previously described^34^.

For immunohistochemical staining, slides were incubated in 5% (w/v) bovine serum albumin (Sigma), 0.5% goat serum (R&D Systems), and 0.1% Triton-X-100 (Amresco) in PBS for 30 minutes. Primary and secondary antibodies were applied for 1 hour and 30 minutes, respectively, followed by Hoechst (Invitrogen) staining to visualize nuclei with PBS washed between staining. The slides were then mounted with Anti-fade mounting medium (Vectashield). The staining included the primary antibodies of laminin (Sigma), embryonic myosin heavy chain (eMHC, DSHB), collagen I (Abcam), and perilipin (Cell Signaling Technology) to assess cross-sectional area, early myofiber regeneration, fibrous infiltration, and fatty infiltration, respectively. Secondary antibodies included Goat anti-mouse IgG H+L AlexaFluor 647 (ThermoFisher) and Goat anti-rabbit IgG H+L AlexaFluor 555 (ThermoFisher). Whole sections were visualized and imaged (Zeiss LSM 700 confocal microscope, 10x) as tile scans (Zen Black, Zeiss) and exported as maximum intensity projections. For each muscle, two sections were evaluated in the MTJ and two in the MB using FIJI ImageJ software (NIH). The “Analyze Particles” tool was used to measure the cross-sectional area of the muscle fibers outlined by laminin-staining. At least 500 randomly selected fibers were assessed, and a random number generator was used to select 500 fibers for each animal. Centrally located nuclei (CLN), eMHC, collagen (fibrosis) and perilipin were normalized by the analysis area.

### 2.9 Statistical Analysis

Error bars represent mean ± standard deviation. Differences between groups were assessed by T-test or 2-way ANOVA with repeated measures where appropriate followed by Tukey’s or Sidak’s *post hoc* test (p<0.05). All analysis was performed using Prism9 (Graphpad).

## 3. RESULTS

### 3.1 Hydrogel Fragment Fabrication and Characterization

The fragmentation process (Fig 1A) reliably produced fragments which upon rehydration after lyophilization provided a controlled injection with a consistency similar to a paste (Fig 1B). This consistency allowed for controlled, localized placement of injections. The fragments had overlapping minimum and maximum lengths (Fig 1C-D) and the average lengths of the fragments from the three batches was 83.22 ± 28.87, 87.99 ± 30.35, and 85.72 ± 33.10 µm, respectively, with no differences between the batches (Fig 1E).

**Figure 1:**
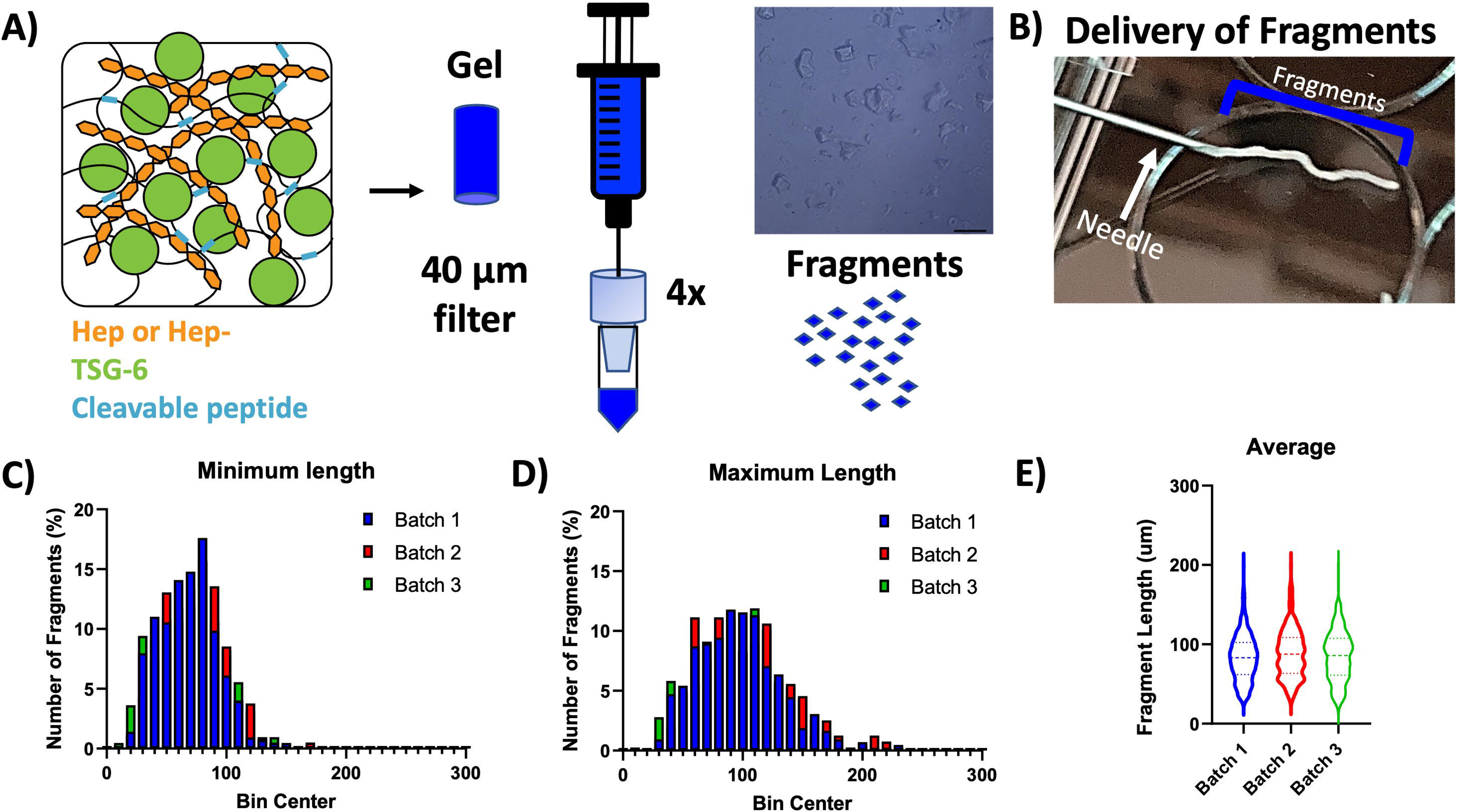
Fragment System and Characterization. **The A)** schematic displays the hydrogel fragmentation process leading to an injectable system **B)** that can be delivered in a controlled manner with **C)** minimum, **D)** maximum and **E)** average lengths of the rectangular fragments displayed. (n=3 batches, p<0.05).

### 3.2 *In vivo* Growth Factor Release

The release of TSG-6 from Hep and Hep-fragment systems can be visualized with decreased signal over the course of the study (Fig 2A) with little to no visible signal on day 21 in both groups. The two groups provided similar release over 21 days with no differences between the groups at any timepoint. The Hep-system demonstrated sustained release throughout the 21 day study and the Hep system showed sustained release through day 14 (Fig 2B).

**Figure 2:**
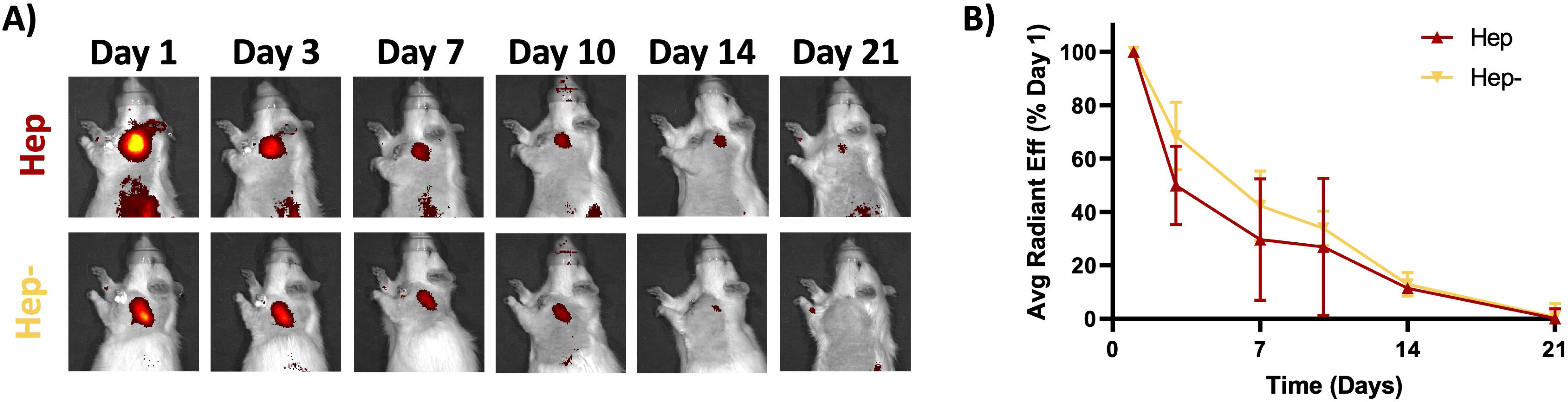
Hep and Hep-produce similar in vivo release profiles. Fragment delivery systems containing either fully sulfated heparin (Hep) or fully desulfated heparin (Hep-) released TSG-6 over the 21-day time period with no differences in release between the two groups at any timepoint (Two-way ANOVA with post hoc test, n=4/grp, p<0.05).

### 3.3 Flow Cytometry

In the MTJ region, flow cytometry (Fig 3A) revealed no differences between groups or over time in T-cells and helper T-cells. However, all groups had a significant increase in regulatory T-cells from Day 3 to Day 7. The Saline group was the only group to show a significant increase in cytotoxic T-cells over time.

**Figure 3:**
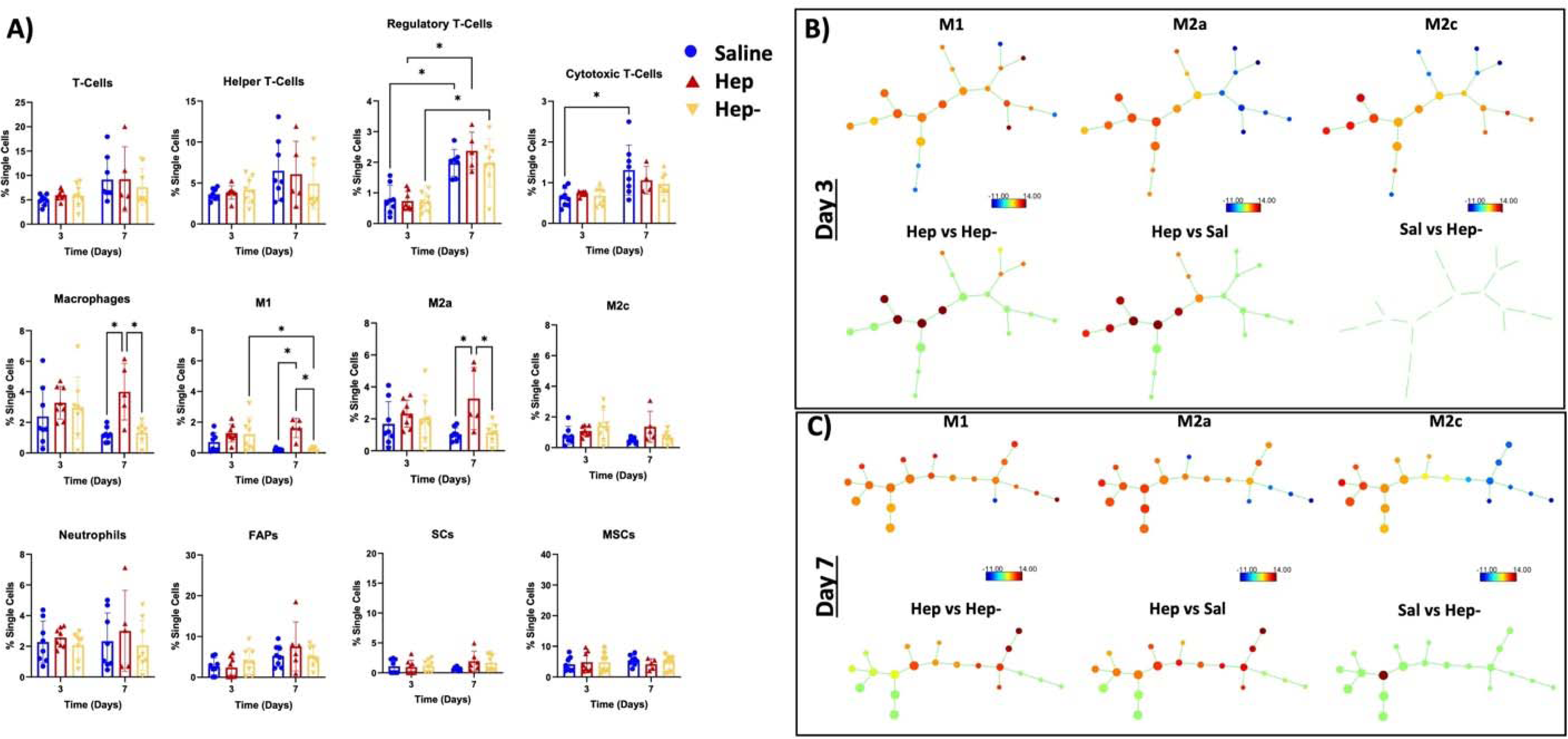
The MTJ region has increased immunomodulation between groups. **A)** The data for each cell type 3 and 7 days after injury with saline (blue), Hep + TSG-6 (red) or Hep-+ TSG-6 (yellow). **B-C)** SPADE analysis of macrophage subtypes showing study population level M1, M2c, and M2a trees (first row of subfigure) and comparisons between groups (second row of subfigure) at **B)** day 3 and **C)** day 7. (Two-way ANOVA with post hoc test, n=5-8/grp, *p<0.05)

In terms of innate inflammatory cells in the MTJ region, there were no differences between groups on Day 3. Interestingly, the total amount of macrophages was significantly higher in the Hep group on Day 7 compared to the Saline and Hep-groups. Specifically, M1 and M2a macrophage subtypes were found to be significantly higher in the Hep group compared to the other two groups on Day 7, whereas all groups had similar M2c macrophages. The Hep-group showed reduced M1 macrophages from Day 3 to Day 7. There were no differences in the M2a or M2c populations over time for this group. There were no differences between groups or over time in neutrophils. In terms of stem cells, no differences were observed between groups or over time for FAPs, SCs or MSCs.

The SPADE analysis (Fig 3B and Fig 3C), which displays the nodes and spatial pattern for each macrophage subtype (top rows), showed clear differences between the Hep treatment and Hep- and Saline (bottom rows) on both Days 3 and 7. Additionally, the changes between those groups overlapped with both M2 and M1 SPADE nodes. The SPADE comparisons between the Hep- and Saline group showed no differences on Day 3 and differences in one node by Day 7.

In the MB region, the flow cytometry analysis (Fig 4A) revealed no differences between groups and time points other than an increase in regulatory T-cells from Day 3 to Day 7 in the Hep-group. SPADE analysis of the MB region (Fig 4B-C) displayed differences in one node in the Hep and Hep-groups at both time points and showed moderate difference between the Hep and Saline groups on Day 3.

**Figure 4:**
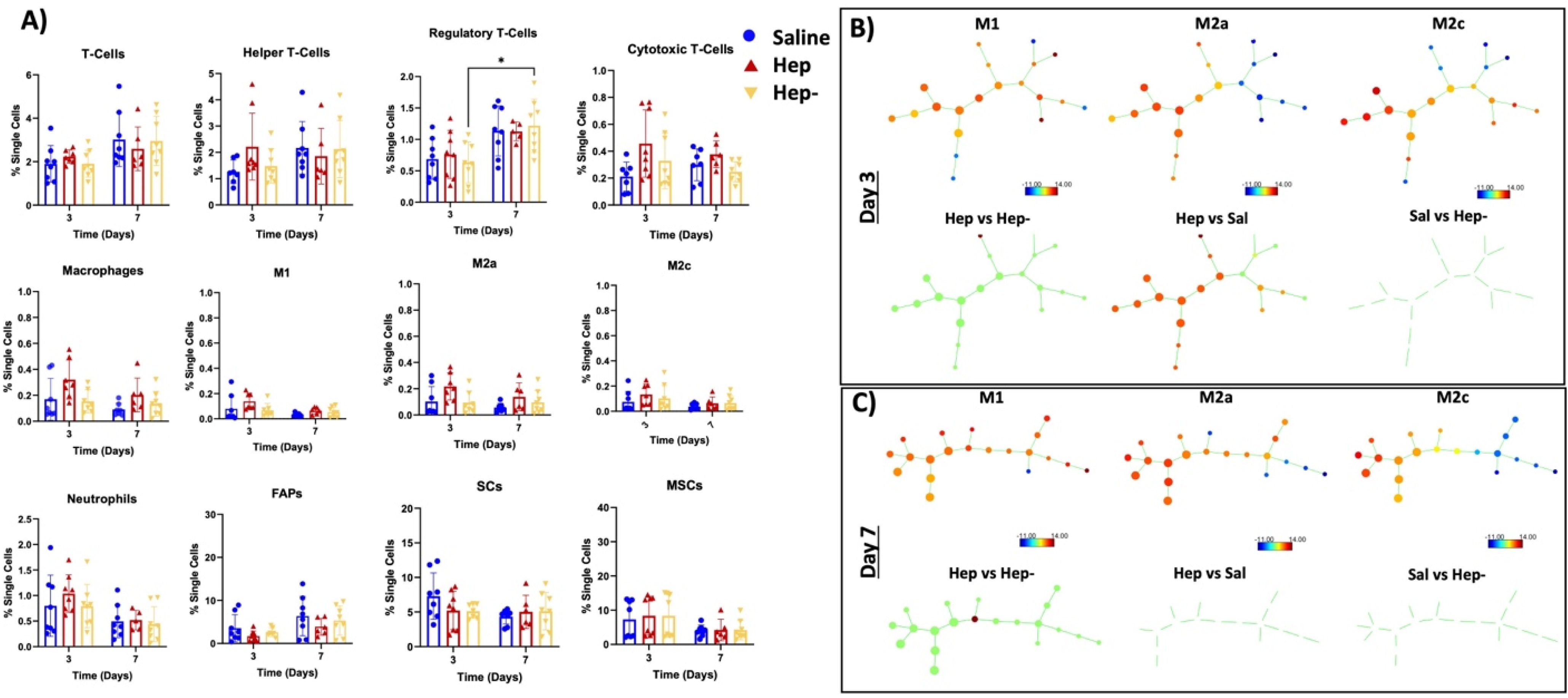
The MB region has largely similar immunomodulation between groups. **A)** The data for each cell type 3 and 7 days after injury with saline (blue), Hep + TSG-6 (red) or Hep-+ TSG-6 (yellow). **B-C)** SPADE analysis of macrophage subtypes showing study population level M1, M2c, and M2a trees (first row of subfigure) and comparisons between groups (second row of subfigure) at **B)** day 3 and **C)** day 7. (Two-way ANOVA with post hoc test, n=5-8/grp, *p<0.05)

### 3.4 Immunohistochemistry

In the MTJ region, the Hep group showed significantly more eMHC fibers than the Saline group on Day 7 (Fig 5A). eMHC staining in the Hep group significantly decreased from Day 7 to Day 14 (Fig 5A). In the MB region, there were no differences in eMHC fibers over time or between groups (Fig 5B).

**Figure 5:**
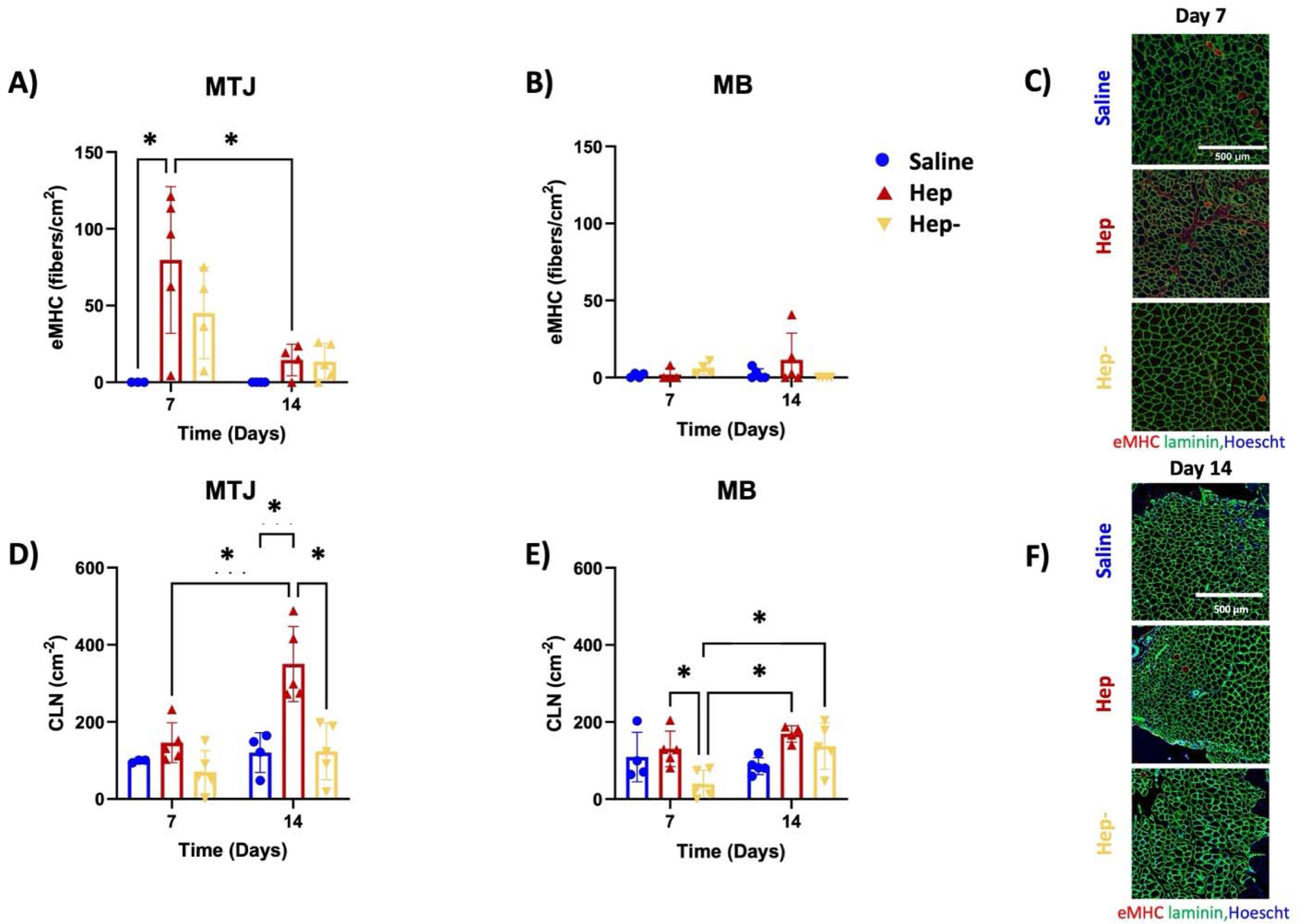
Hep + TSG-6 produces an increase in metrics of muscle regeneration (eMHC staining and number of centrally located nuclei). eMHC of **A)** MTJ and **B)** MB, and centrally located nuclei (CLN) regenerating fibers measured as number of positive fibers per area in the **D)** MTJ and **E)** MB. **C)** and **F)** Representative cross-sectional area (laminin, green), eMHC positive (red) fiber, and Hoescht (blue). (One or two-way ANOVA with post hoc test, n=3-5/grp, *p<0.05)

In the MTJ region, the Hep group showed significantly more CLN than the Hep- and Saline groups on Day 14 and demonstrated an increase from Day 7 to Day 14 (Fig 5D). In the MB region, the Hep group demonstrated significantly more CLN than the Hep-group on Day 7 and only the Hep-group showed increased CLN from Day 7 to Day 14 (Fig 5E).

In the MTJ region on Day 7, the Hep and Saline groups showed significantly greater number of small fibers (0-200 µm^2^) than the Hep-group, and the Hep and Hep-groups had greater 200-400 µm^2^ small fibers than the Saline group (Fig 6A). The Hep and Hep-groups contained significantly greater number of small fibers than the Saline group in the MB on Day 7 (Fig 6B). On Day 14, the Hep and Hep-groups showed significantly greater number of small fibers and fewer medium sized fibers (1000-1200 µm^2^) than the Saline group in both the MTJ and MB regions (Fig 6C-D).

**Figure 6:**
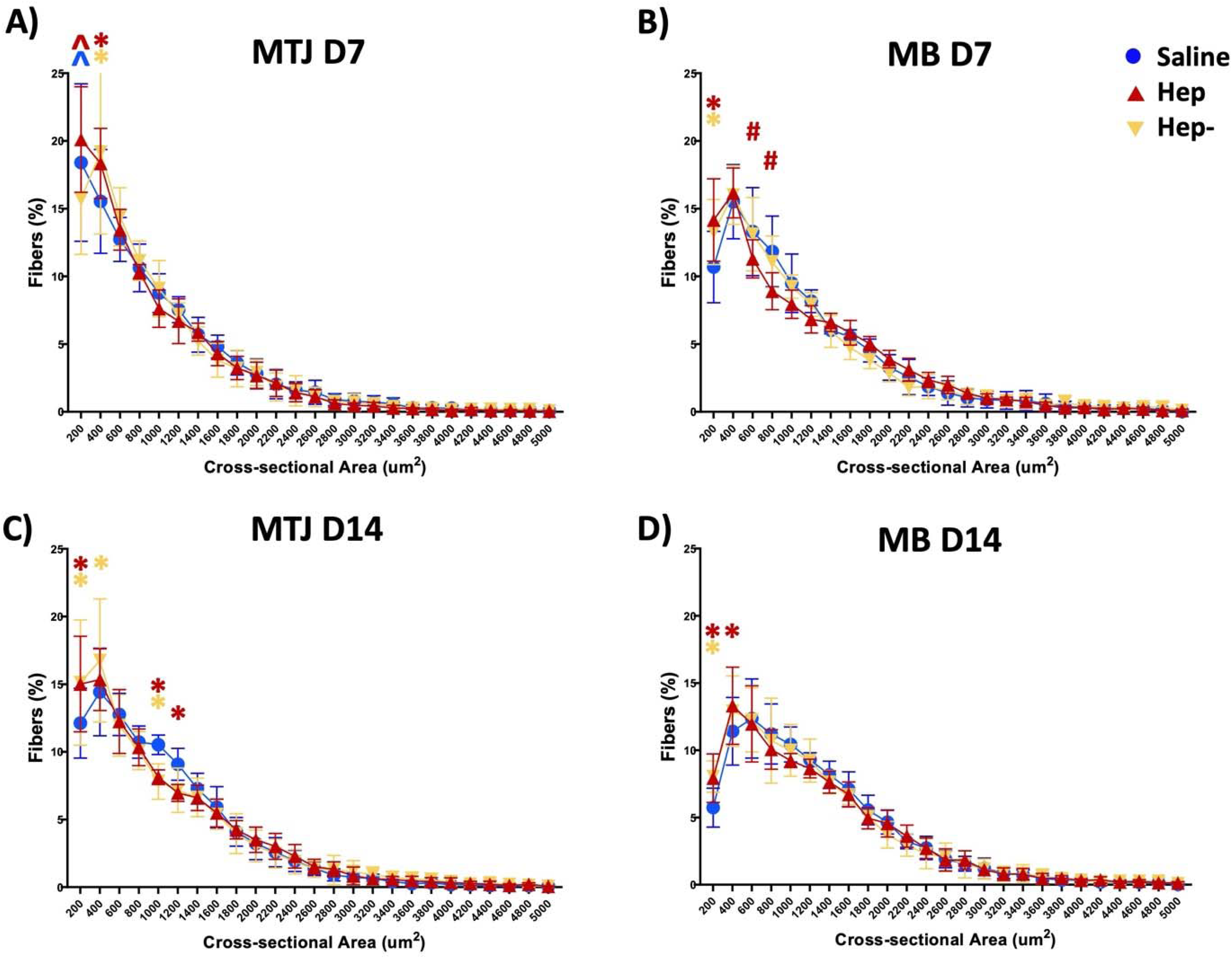
Hep + TSG-6 produces an increase in small fibers. Cross-sectional area of 500 random fibers from each group at. **A)** MTJ and **B)** MB at day 7 with and **C)** MTJ and **D)** MB at day 14. (Two-way ANOVA with post hoc test, n=4-5/grp, p<0.05, *different from saline, ^group of that color vs. all other groups, # different from Hep- and from Saline group)

## 4. DISCUSSION

In this work, a degradable PEG-based granular hydrogel system containing two heparin derivatives was developed with sustained release profiles *in vivo* to understand the interactions of TSG-6 and the heparin derivatives. The cellular and regenerative responses differed between the three *in vivo* groups (Saline, Hep and Hep-). In particular, the Hep group had heightened macrophage response over time and increased markers of muscle regeneration (eMHC and CLN on day 7 and 14, respectively). In addition to the release system findings, we discovered that, similar to other studies^34^, the MTJ region has an increased level of regeneration-associated activity and cellular changes, whereas the injury and treatments showed less differences in the MB region.

As displayed in Figure 1a, our injectable hydrogel-based system allows for formation of reproducible fragments and high concentration injections with spatial control, aiding in repeatability and reliability in release (Fig 2). Granular hydrogels have been manufactured via a variety of methods including microfluidic devices, batch emulsion, and extrusion fragmentation followed by jamming. The manufacturing method can change the hydrogel properties including size and porosity.^35,36^ Granular hydrogels can also be adapted, pre-swollen and added with fibrous structures to change the mechanical properties of the overall structure.^37^ Microgels have previously been annealed via various chemistries after microgel formation and drying processes to form a porous scaffold-like structure, better known as MAPs.^38–41^ Heparin has also been included in MAPs to encourage tissue infiltration through sequestration of growth factors.^42^

In contrast, the microgel fragments formed in this study did not have chemical post-processing leading to an annealed scaffold-like structure. The fragments were, however, lyophilized which may have induced some interactions between fragments.^41^ Additionally, the various fragment shapes may have caused some fragment-to-fragment interaction. Furthermore, the density of the muscle tissue and injection along the scapular spine may have enhanced the repeatability and retention of the fragments. For other tissues, annealing or other means of particle bonding may be crucial for repeatable placement during injection.

In order to select the heparin derivatives tested in this study, an *in vitro* screen was conducted to demonstrate the effects of the various GAGs on TSG-6 activity over time (Fig S1). In this assay, heparin should potentiate the actions of TSG-6 with its co-factor inter-alpha-inhibitor (IαI) in inhibiting plasmin (black bar Figure S1B)^18^. Based on these results, the fully sulfated heparin derivative (Hep) was the only derivative to show this increased anti-plasmin activity at 50 min and maintain this activity, to some degree, up to 48 hours. Other derivatives did not demonstrate a reduction in plasmin activity (beyond that seen with TSG-6 + IαI) and performed similarly to each other, so the fully desulfated molecule (Hep-) was chosen as a control for these studies. Prior to *in vivo* release, we also performed *in vitro* release studies with PDGF (pI 9.8)^43^, with a similar charge as TSG-6 (pI 9.48)^44^, to ascertain Hep or Hep-to TSG-6 molar ratios that produce similar release profiles (Fig S2). Using the information gained from these experiments, we then selected formulations for *in vivo* release studies. The *in vivo* release of TSG-6 was ultimately found to be similar between the Hep and Hep-groups (Fig 3B), which allowed for the evaluation of differences resulting from the interaction of TSG-6 with Hep and Hep-.

The flow cytometry results on Day 3 and Day 7 (Fig 3A and 4A), demonstrate that, overall, there are more changes in cell populations in the MTJ than MB regions after injury and treatment, consistent with our previous work^34^. In terms of acquired immunity, increased levels of regulatory T cells were observed in all groups in the MTJ from Day 3 to Day 7 and in cytotoxic T-cells in the Saline group only in both regions. Overall, the increase in T-regs over time suggest a healing response from each group following injury with increased T-regs generally associated with the transition from an inflammatory to anti-inflammatory environment.^45^ Generally, an increase in cytotoxic T-cells is associated with the inflammatory stage of the injury response. ^45^ These results suggest the Hep and Hep-groups may be progressing towards T-cell population indicative of resolution of acute inflammation earlier than the Saline group.

Supporting this conclusion, comparison of the flow cytometry results for macrophage subpopulations in each muscle region as indicated by the SPADE analysis (Fig 3B-C and 4B-C) demonstrated distinctions in macrophage sub-populations over time based on treatment type. SPADE analysis is a cellular clustering tool used to generate dendrograms of pooled cellular data into nodes along a trajectory to track lineages and cellular transitions^46^. The M1, M2c, and M2a SPADE trees (top rows Fig 3B-C and 4B-C) show that fewer right-sided nodes are associated with the M2c and M2a population (green to blue nodes), whereas M1-like macrophages are more associated with the right-sided nodes (more yellow to red nodes; particularly obvious on Fig 3C and 4C). The left side nodes represent populations from all subtypes, as displayed by the M1, M2c, and M2c trees (yellow to red nodes in left of all graphs). These differences suggest that a decrease in right-sided node populations combined with an increase in left sided node populations is likely representative of M1 to M2 transition.

With this representation, interrogation of the macrophage populations with SPADE analysis revealed differences between the Hep group and the other groups in all regions and timepoints, with the differences in nodes skewed to the left side of the tree (bottom rows Fig 3B-C and 4B-C), suggesting a transition to a more anti-inflammatory response, despite a relatively higher number of M1 macrophages in this group in the MTJ region at Day 7 (Fig 3A). This interpretation is supported by the simultaneous demonstration of a higher number of M2a macrophages compared to other groups in the MTJ region at Day 7 (Fig 3A). In a similar vein, the Hep-group showed distinctions from the Saline group in the SPADE analysis on Day 7 in the MTJ region, with decreases at all nodes (green) except for one left sided node (Fig 3C). Such analysis supports that the Hep-is trending more towards an anti-inflammatory milieu compared to the Saline group, and this is corroborated by the significant decrease in M1 macrophages over time in the MTJ region (Fig 3A)

Taken together, the data suggest that Hep group demonstrates a prolonged anti-inflammatory effect with an overall increased macrophage response compared to no treatment, with a more muted response by the Hep-group. Evidence shows that macrophages have the plasticity to transition between types in response the environment. ^47^ As previously mentioned, TSG-6 is known to contribute to M1 to M2 transition^11^, so the macrophage phenotypic shifts observed may be a result of heparin-related potentiation of TSG-6^18^ or prolongation of its bioactivity^20,48^. The potentiation or prolonging of the effects of TSG-6 by fully sulfated heparin may be due to the binding interaction of heparin to TSG-6. It has been postulated that the 2-O and 6-O-sulfates are prominent in the interaction of heparin with TSG-6^49^, so the desulfated heparin derivative may have less interaction with TSG-6, thus leading to reduced effect on the cellular milieu. It should be noted that, while we have shown that there is little additional inflammatory effect of a similar hydrogel-based system injected in the same animal model^25^, further studies would be required to fully differentiate the effects of the carrier vs TSG-6 on the inflammatory environment in this injury model.

Interestingly, our study did not reveal any differences in neutrophil, MSC, SC, or FAP populations. TSG-6 is known to affect the migration of neutrophils through hyaluronan, inter-alpha inhibitor interaction, and direct interaction with chemokine CXCL8.^50,51^ The change in migration of neutrophils has been linked with reduced MMP9 in a brain injury model with TSG-6 treatment.^52^ Since no differences were found between groups, it is likely that the delivery of TSG-6 in combination with Hep or Hep-did not change these interactions in the rotator cuff tear environment.

The immunostaining results (Figures 5-6) support an increased level of regeneration in the Hep group, with significantly higher eMHC on Day 7 followed by significantly higher CLN on Day 14 compared to other groups. In addition, the Hep group showed an increased number of smaller fibers than saline at both timepoints (Fig 6), which can be indicative of regenerating fibers^53,54^. eMHC staining and more centrally-located nuclei are indicative of early and progressing muscle regeneration,^4,55–57^ leading to myofibril and eventually myofiber formation. Various cell types, including T-cells, macrophages, and associated subtypes progress from inflammatory to anti-inflammatory leading to muscle regeneration.^45^ In particular, pro-inflammatory macrophages encourage activation and proliferation of satellite cells in response to injury, whereas anti-inflammatory macrophages encourage differentiation to form myofibers.^58,59^. Thus, the combination of the increased and prolonged macrophage response and increase in regulatory T cells over time observed in the Hep group suggests that the combination of heparin and TSG-6 may act to change the local cellular milieu to one that encourages early muscle regeneration in this model.

In parallel with exploring how release of TSG-6 from heparin gels affects local cellular response, we also investigated the potential of this system to reduce plasmin and thus MMP activity *in vivo* ^18,60^ as another means to limit muscle damage after rotator cuff tendon tear (Fig S3). We found that MMP 2/9 activity actually increased in the Hep group on Day 7 compared to the Saline group. This heightened MMP activity combined with no differences in neutrophils suggest that these results may be driven by infiltration of macrophages, which secrete a variety of MMPs as a part of the acute inflammatory response^61^ that may overwhelm or circumvent the capacity of TSG-6/heparin to reduce plasmin activity. Muscle repair requires tissue remodeling and thus early MMP activity may be necessary^62,63^. It should be noted there is no difference in MMP activity between groups by Day 14, which correlates with increases in CLN (a later marker of muscle regeneration) in the heparin group.

A variety of other studies have investigated the delivery of soluble TSG-6 or TSG-6 secreting cells.^8,64^ Similar to our findings, many have found that TSG-6 encouraged macrophages towards anti-inflammatory polarization or reduced inflammation including in an inflammatory lung injury,^11^ colitis,^12^ alcohol hepatitis^13^, and stroke.^65^ These studies suggested that this transition was regulated through the TSG-6 effect on SOCS3, STAT3, STAT1, oxidative stress, and NFkB. Other studies have found that TSG-6 decreased inflammatory cytokines which led to reduced scar formation in a rabbit ear model^66^ and early gingival wound healing^67^. Although these studies produced similar findings regarding inflammation, none utilized a delivery platform. Previous work in our laboratory compared short-term release of TSG-6 from an N-desulfated heparin (Hep^-N^) microparticle system to a soluble TSG-6 injection in a rat medial meniscal transection model of osteoarthritis^18^. It was found that that the Hep^-N^ TSG-6 group reduced cartilage damage compared to triple the dose of soluble TSG-6. This work expanded on our previous platform by comparing two heparin derivatives demonstrating sustained release over 14+ days, but similarly found that a more highly sulfated carrier resulted in less damage/more metrics of tissue regeneration.

It should be noted that, in this work, treatment was administered immediately after tendon injury, which is not how patients typically present^68–70^, and that the animal model used did not include tendon reattachment, which is the current standard of care for massive rotator cuff tendon tear^71,72^. Thus, while the findings herein provide initial data supporting muscle treatment with TSG-6 after rotator cuff tear, future work should include examining the effects of a delayed treatment strategy in animal models that include tendon reattachment.

## 5. CONCLUSIONS

In these studies, we developed a PEG-based injectable granular hydrogel containing two heparin derivatives (Hep and Hep-) as well as an MMP-degradable peptide to promote sustained release of TSG-6 over 14+ days *in vivo* in a rat model of rotator cuff tear. The system was designed to demonstrate similar release profiles, thus facilitating comparisons between delivery from different heparin derivatives on the local cellular milieu and level of tissue repair in two different areas of muscle (near the myotendious junction and further into the muscle belly) that been shown previously to have differing responses to rotator cuff tendon injury^34^. Results showed controlled delivery of TSG-6 from the fully sulfated heparin group caused heightened macrophage response over time and increased markers of early muscle regeneration (eMHC and CLN on day 7 and 14, respectively), particularly in the MTJ region of the muscle, compared to release from desulfated heparin hydrogels. In addition to providing a strategy for localized, controlled delivery of TSG-6 to muscle, these results demonstrate the potential of key pairings of biomaterial substrates and cargo to promote tissue healing.

## ACKNOWLEDGEMENTS

The authors would like to acknowledge Dr. Lauren Hymel and Dr. Thomas Turner (Botchwey Laboratory, Georgia Tech) and Dr. Molly Ogle (Temenoff Laboratory, Georgia Tech), for guidance on flow cytometry, SPADE and associated analysis, and immunohistochemical staining and analysis.

## AUTHORSHIP CONTRIBUTION STATEMENT

**<u>Joseph Pearson:</u>** Conceptualization, Data Curation, Formal Analysis, Methodology, writing – original draft

**Jiahui Mao:** Data Curation, Formal Analysis, Writing – review and editing

**Johnna Temenoff:** Conceptualization, Funding acquisition, Project administration, Supervision, Writing – review and editing

## DISCLAIMER

The content is solely the responsibility of the authors and does not necessarily represent the official views of the National Institutes of Health.

