## Supplemental Data for "Effects of Release of TSG-6 from Heparin Hydrogels on Supraspinatus Muscle Regeneration"

**SUPPLEMENTAL INFORMATION**

**S1. Supplemental Methods:**

***S1.1 In vitro* Plasmin Inhibition Assay:**

*In vitro* plasmin inhibition studies were completed as previously reported.^1^ Briefly, 0.5% wt bovine serum albumin blocking buffer was added to 96-well plate for 1 hour. TSG-6 was added in a 7.4 pH buffer solution of 10 mM HEPES (Sigma), 150 mM NaCl, and 0.02% v/v Tween 20 (VWR) in dH_2_O at 108 nM TSG-6 alone (blue bar, Fig S1) or with Hep, Hep- or N-desulfated heparin (Hep^-N^) and incubated for 24 hours at 37°C. Then inter-alpha inhibitor (Athens Research & Technology, 24 nM) was added and allowed to incubate for 30 minutes at 37°C. Lastly 3.4 nM plasmin (Sigma) and 197 µM plasmin substrate (N-p-tosyl-Gly-Pro-Lys 4-nitroanilide acetate salt, Sigma) were added and incubated at room temperature for 20 minutes and 37°C for 20 minutes, at which time the plate was analyzed on a plate reader (Biotek Synergy H4,) at 405 nm at 50 minutes (24 hours + 50 minutes) and 24 hours (total 48 hours). The Hep^-N^ in this study was synthesized similarly to Hep- other than the heparin pyridinium was dissolved at 1 mg mL−1 in 9 : 1 v/v dimethyl sulfoxide (DMSO)/dH2O at 50 °C for 2 hours.^1^

***S1.2 In vitro* release study:**

For *in vitro* release studies, a model protein (PDGF-BB (Preprotech), 1 µg, at different molar ratios to heparin) was fluorescently tagged (Dylight 488) in a similar fashion to the IVIS study and was loaded onto the Hep or Hep- fragments at molar ratios of 1:1, 5:1, 10:1 and 100:1, 1000:1, respectively (PDGF-BB:Heparin). A MMP control group without heparin was also evaluated. Release (n=4/group) was conducted in 0.1% bovine serum albumin solution (Fisher, dissolved in PBS) and quantified by fluorescence plate reader at 528 nm in the release medium over 5 days.

***S1.3 In vivo* MMP Assay:**

Muscles were excised following euthanasia, finely minced, placed in lysis buffer consisting of 20mM Tris–HCl (Amresco), 5mM EGTA (Sigma), 150mM NaCl (Fishersci), 20mM glycerol-phosphate (Alfa Aesar), 10mM NaF (Sigma), 1mM sodium orthovanadate (Sigma), 1% Triton-X 100 (Amresco), and 0.1% Tween 20 in 500 mL deionized water and 0.1 mM of fresh leupeptin (Fisher), and then ground using sample grinding kit (Cytiva).^2^ The samples were evaluated for total protein using a Bicinchoninic acid (BCA) assay (Thermofisher). MMP activity was measured using MMP 2/9 Fluorogenic Assay (Sigma). Briefly, the samples were diluted in the assay activation buffer and combined with the substrate working solution and incubated at 37°C for 6 hours at which time the plate fluorescence was read with emission/excitation of 320/405. The MMP activity is shown as MMP/protein (n=5-6/group).

**S2. Supplemental Results:**

***S2.1 In vitro* Plasmin Inhibition Assay:**

The plasmin inhibition at 50 minutes in the Hep group was significantly greater than the TSG-6, IaI alone, and plasmin control groups, and the Hep- and Hep^-N^ groups had significantly greater inhibition than the IaI alone and plasmin control groups but not the TSG-6 group (Fig S1). The plasmin inhibition at 48 hours in the Hep group was significantly greater than the IaI alone and plasmin control groups, and the Hep- and Hep-N groups had significantly greater inhibition than only the plasmin control group.

***S2.2 In vitro* release study:**

For *in vitro* release studies, all ratios of the Hep and Hep- groups had similar release over time. The Hep 10:1 group showed significantly lower release at days 3 and 5 compared to the MMP control.

***S1.3 In vivo* MMP Assay:**

The MMP activity in the Hep group was significantly greater than the saline and Hep- groups on day 7. There were no differences in MMP activity between groups on day 14.

**FIGURES**

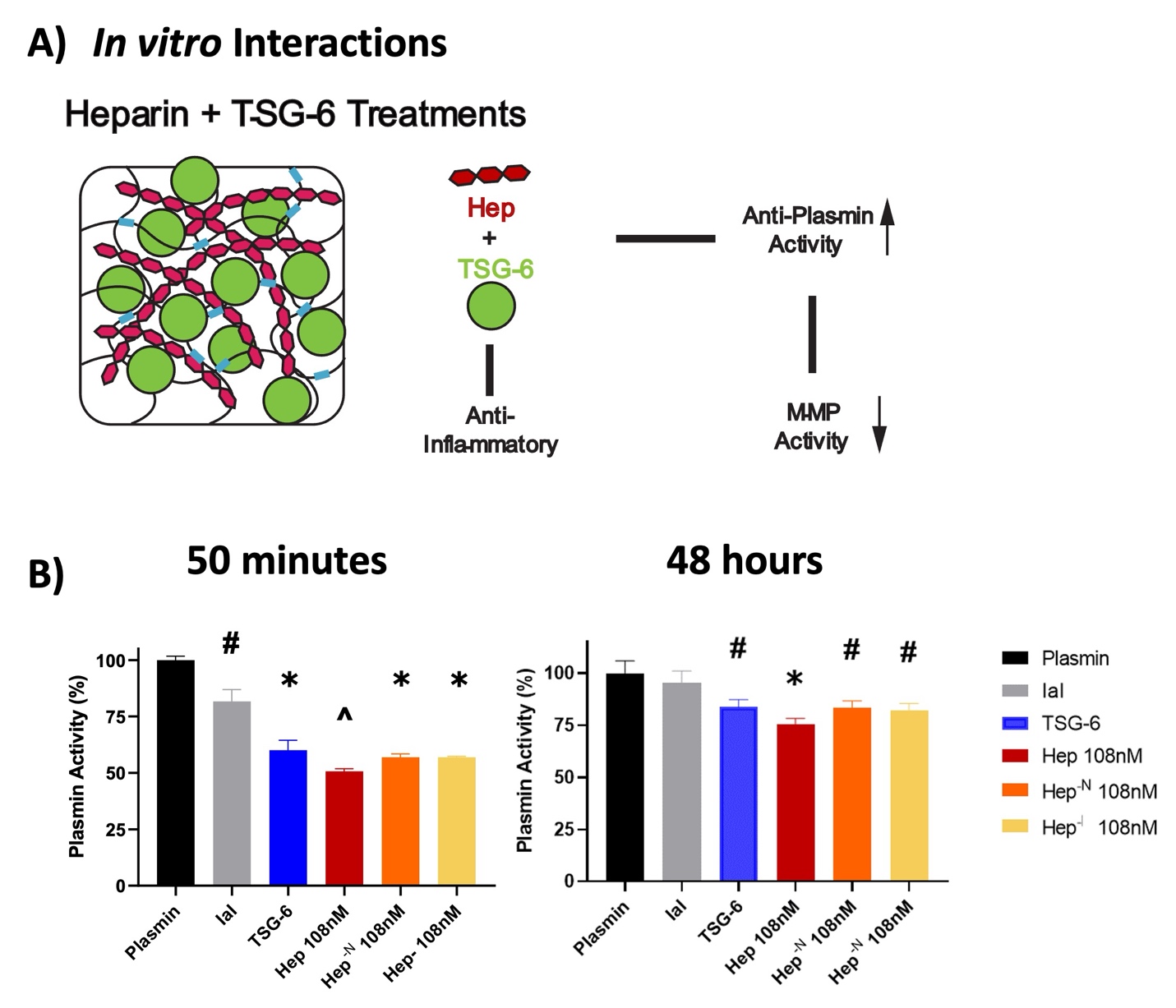

**Figure S1: TSG-6, heparin interaction reduces plasmin activity in vitro. A)** Depiction of hypothesis for these studies: Fully sulfated heparin increases TSG-6 anti-plasmin activity; plasmin aids in MMP activation, so fully sulfated heparin may lead to decreased MMP activity. **B)** Anti-plasmin activity is potentiated by fully sulfated Hep as the only heparin derivative significantly different from TSG-6 alone at 50 minutes and IaI after 48 hours. (One-way ANOVA with posthoc test, n=4/grp, p<0.05, significance from #Plasmin, *IaI and Plasmin, and ^TSG-6, IaI and Plasmin).

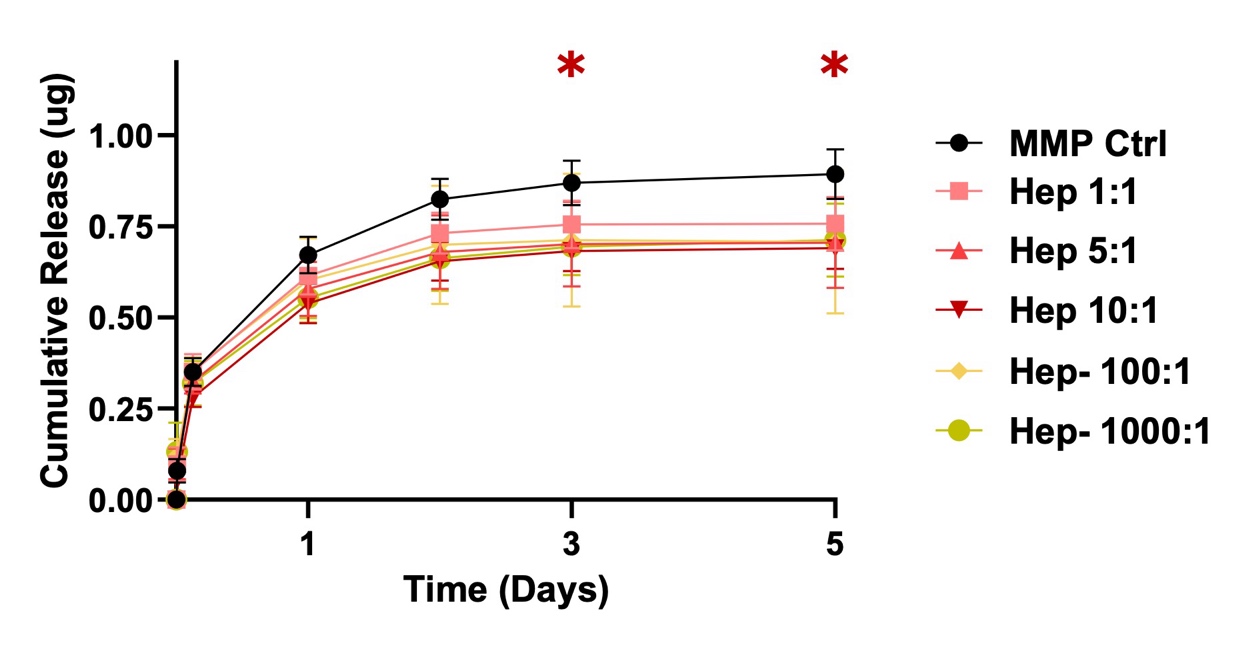

**Fig S2: Changing protein: heparin molar ratios produce similar release profiles of PDGF-BB (model protein).** Fluorescently tagged PDGF-BB (1 µg) released from Hep or Hep- containing fragments over 5 days with varying Hep:PDGF-BB molar ratios. (Two-way ANOVA with post hoc test, n=4/grp, *color of group significantly greater than the MMP Ctrl at that time point, p<0.05).

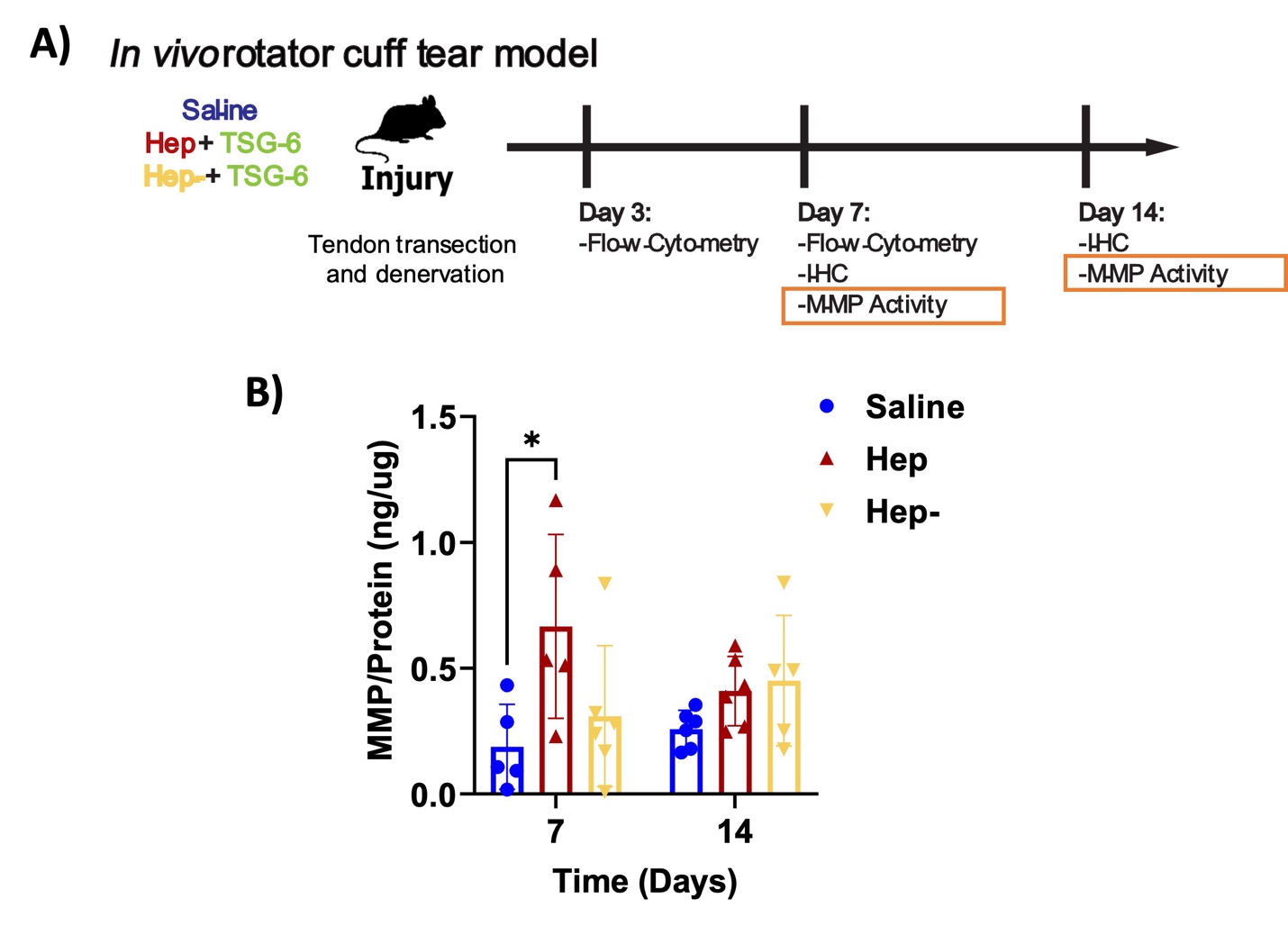

**Figure S3: TSG-6 - heparin group shows increased MMP 2/9 activity in vivo at early timepoints. A)** In vivo study design to examine **B)** MMP activity after treatment (Two-way ANOVA with post hoc test, n=3-6/grp, *p<0.05.).
